# Left-right asymmetry is formed in the basal bodies of the mouse node cilia in a cilia motility-dependent manner

**DOI:** 10.1101/2023.09.13.557556

**Authors:** Hiroshi Yoke, Atsushi Taniguchi, Shigenori Nonaka

**Affiliations:** Laboratory for Spatiotemporal Regulations, National Institute for Basic Biology (NIBB), Okazaki 444-8585, Japan; Present address: Research Center of Mathematics for Social Creativity Research Institute for Electronic Science Hokkaido University, Sapporo 001-0020, Japan; Spatiotemporal Regulations Group, The Exploratory Research Center on Life and Living Systems (ExCELLS), Okazaki 444-8585, Japan

## Abstract

Laterality of the shapes and arrangements of the visceral organs in mice is determined in the node, a small cavity found at the ventral side of 7.5 dpc (days post coitum) embryos. On the node cells, motile cilia which are tilted toward the posterior side of the embryos show clockwise movement and thus produce fluid flow in the node toward the left side of the embryos. This left-ward flow regulates left/right (L/R) asymmetric gene expressions and L/R asymmetric morphogenesis in later stages. Structurally, node cilia have the characteristics of primary cilia and their basal body (mother centriole) is accompanied by a daughter centriole. Here, to obtain insights into the process of symmetry breaking by node cilia, we investigated whether the structure of the cilia themselves have L/R asymmetry, and found that positions of the daughter centrioles become biased to the right side of the mother centrioles in a stage-dependent manner. We found that this L/R asymmetry of the basal bodies is absent in *iv* mutant mice, in which node cilia are immotile, suggesting that formation of this L/R asymmetry in the basal bodies requires cilia motility. It has been reported that culturing embryos in a flow chamber with artificial counter-flow, which is toward the opposite direction to the endogenous leftward flow in the node, results in reversed laterality of the visceral organs in later stages. However, we found that applying such artificial counter-flow did not reverse the L/R asymmetry of the basal bodies, and the daughter centrioles were still biased to the right side of the mother centrioles, suggesting that the L/R asymmetry of the basal bodies is formed independently from the direction of the fluid flow in the node and that it is independent from the laterality of the visceral organs. Although the biological significance of this phenomenon is unknown so far, these results suggest that node cilia have a previously unknown mechanism to produce L/R asymmetry in the basal bodies inside the cells in early development, independently from the canonical fluid flow-dependent L/R determining pathway.

## Introduction

The body plans of the animals have L/R asymmetry including the shape of the heart and the arrangements of the guts and other visceral organs. It is considered to be important for packing of the organs in the body and proper functioning of them.

During development of the mouse, the first molecular asymmetry is established in a small cavity called the node at the ventral side of a 7.5 dpc embryo (Sulik et al., 1994; Little and Norris, 2020). On the apical surface of the node cells, primary cilia protrude toward the node cavity with tilting toward the posterior side of the embryos (Nonaka et al., 2005). These posterior-tilted cilia produce directional fluid flow within the node toward the left side of the embryos by showing chiral rotational movement in a clockwise direction (seen from the tip side of the cilia) (Nonaka et al., 2005; Okada et al., 1999; Okada et al., 2005). This leftward fluid flow called a “nodal flow” regulates L/R asymmetric expression of the genes such as *Nodal* and *Pitx2* and L/R asymmetric formation of the visceral organs such as the heart (Lowe et al., 1996; Nonaka et al., 2002). This mechanism of producing L/R asymmetry by node cilia via the direction of the fluid flow is conserved among some vertebrates including mice, rabbits, frogs (Xenopus) and fish (medaka and zebrafish) (Okada et al., 2005; Schweickert et al., 2007; Essner et al., 2005). In the chick and the pig, however, the ventral side of the left-right organizer (the node or Hensen’s node) is covered by endoderm cells and it seems that there is no space where nodal flow can be produced (Essner et al., 2002; Stephen et al., 2014; Gros et al., 2009). Thus, fluid flow-dependent symmetry breaking may not be the only mechanism of producing L/R asymmetry in vertebrates, but a possibility of the variety in the mechanisms of initial symmetry breaking is poorly understood.

To obtain insights into the mechanism of symmetry breaking by node cilia, we sought to investigate whether the node cilia have structural L/R asymmetry. Mouse node cilia have structural characteristics of primary cilia, and in most of them, the structure of the shaft of the cilia is relatively symmetric, with so called “9+0 structure” without central pair microtubules (Takeda et al., 1999; Odate et al., 2016), which is in contrast to motile cilia in multiciliated cells with “9+2 structure” with a polarity defined by the arrangement of central pair microtubules (Choksi et al., 2014; Robinson et al., 2020). The structure at the base of each cilium, from which a cilium originates, is called a basal body, and it is consisted of a microtubule-based barrel-like structure, a centriole. The basal body of a cilium in multiciliated cells does not possess an accessory centriole but has a polarity defined by the position of a protrusion called a basal foot (Kunimoto et al., 2012; Mitchel et al., 2007; Robinson et al., 2020). In contrast, primary cilia including mouse node cilia have an accessory centriole beside the basal body (Nonaka et al., 1998; Hagiwara et al., 2008). These two centrioles are derived from a centrosome, a pair of centrioles that function at the cell division. In quiescent or interphase cells, the mother centriole of the centrosome functions as the basal body and the primary cilium is nucleated from it, while the daughter (or accessory) centriole sits beside the mother centriole (Sorokin, 1962; Kobayashi and Dynlacht, 2011). However, it has been unknown whether the basal bodies of the node cilia have any structural polarity or L/R asymmetry.

Here we investigated the structural asymmetry of the basal bodies of the node cilia and found that positions of the daughter (accessory) centrioles become biased to the right side of the mother centrioles (basal bodies) in a stage-dependent manner. We found that formation of this L/R asymmetry in the basal bodies requires a cilia motility. Unexpectedly, we found that formation of this L/R asymmetry of the basal bodies was independent from the direction of the fluid flow in the node, suggesting that this phenomenon may represent a novel mechanism of symmetry breaking, in which motility of the node cilia produces L/R asymmetry in the basal bodies inside the cells independently from the fluid flow outside the cells.

## Results

### Daughter centrioles become biased to the right side of the mother centrioles in the mouse node in a stage-dependent manner

In order to obtain insights into the process of symmetry breaking by cilia, we investigated whether the structure of the basal bodies of the node cilia have L/R asymmetry. First, we tested whether the arrangements of the basal feet around the basal bodies of the node cilia have L/R asymmetry. In case of cilia in multiciliated cells, each basal body has a single protrusion called a basal foot, and the position of the basal foot defines the polarity of the cilia and is crucial for coordinated ciliary beating (Kunimoto et al., 2012). We found, however, centriolin, which localizes to basal feet, showed ring-like localization without apparent L/R asymmetry around the basal body in the node (Fig. S1). This suggests that multiple of basal feet surround the basal body in the node cells unlike in multiciliated cells. This is consistent with reports in primary cilia in other tissues (Hagiwara et al., 2008; Fewell and Dutcher, 2020).

Then we tested whether the relative positions of mother centrioles and daughter centrioles have L/R asymmetry. As a marker for mother centrioles in immunostaining, we chose a protein called ninein, which localizes to basal feet on the mother centrioles. By double-staining γ-tubulin, which localizes to both mother and daughter centrioles, and ninein, we found that ninein predominantly localizes to one of the pair of centrioles and that the centrioles with ninein signal can be distinguished as mother centrioles in most of the basal bodies (Fig. 1A). Ninein often showed a ring-like structure around the mother centriole (Fig. 1A), probably showing the arrangements of basal feet surrounding the mother centriole, consistent with the result of immunostaining of centriolin (Fig. S1). We first analyzed the positions of the mother centrioles and the daughter centrioles in the node of a late bud embryo. Late bud is a relatively early stage when cilia do not move actively nor produce directional flow yet (Okada et al., 1999). In late bud, early L/R asymmetric signals induced by nodal flow have not yet occurred, such as L/R asymmetric frequencies of the Ca^2+^ spikes at the peripheries of the node (Takao et al., 2013) and degradation of Dand5/Cerl2 mRNA at the left side of the node (Shinohara et al., 2012; Minegishi et al., 2021). Here we categorized the relative positions of the daughter centrioles into two categories: the daughter centrioles at the left side of the mother centrioles and the right side of the mother centrioles (the relative positions as to the anterior-posterior axis is discussed later). We found that in late bud embryos, the numbers of the daughter centrioles at the left side of the mother centrioles and those at the right side of the mother centrioles in the node were comparable, and no significant L/R bias was detected in the 6 embryos investigated (Table 1, Fig. 1B,C). In contrast, we found that the positions of the daughter centrioles were biased to the right side of the mother centrioles in later, somite stages (Table 2, Fig. 1D). In somite stages such as somite-2, rotating cilia produce laminar leftward flow (Okada et al., 1999), and Ca^2+^ signals and Dand5/Cerl2 mRNA have begun to show L/R asymmetry (Takao et al., 2013; Shinohara et al., 2012; Minegishi et al., 2021). We investigated 8 embryos from somite 1–3, and found that in all of them, the daughter centrioles tended to be at the right side of the mother centrioles, and that in 6 of them the bias was statistically significant by the binomial test (*P*<0.05, Table 2). The percentage of the daughter centrioles at the right side of the mother centrioles increased from late bud (50%) to late headfold (LHF) stage (57%) and also from late bud to somite-2 (62%) or to somite-3 (62%) stages (Fig. 1E, Tables 1,2). These results suggest that the daughter centrioles in the node cells become biased to the right side of the mother centrioles in a stage-dependent manner, which is a previously unknown L/R asymmetry formed in the node cells in the early development.

**Fig. 1.**
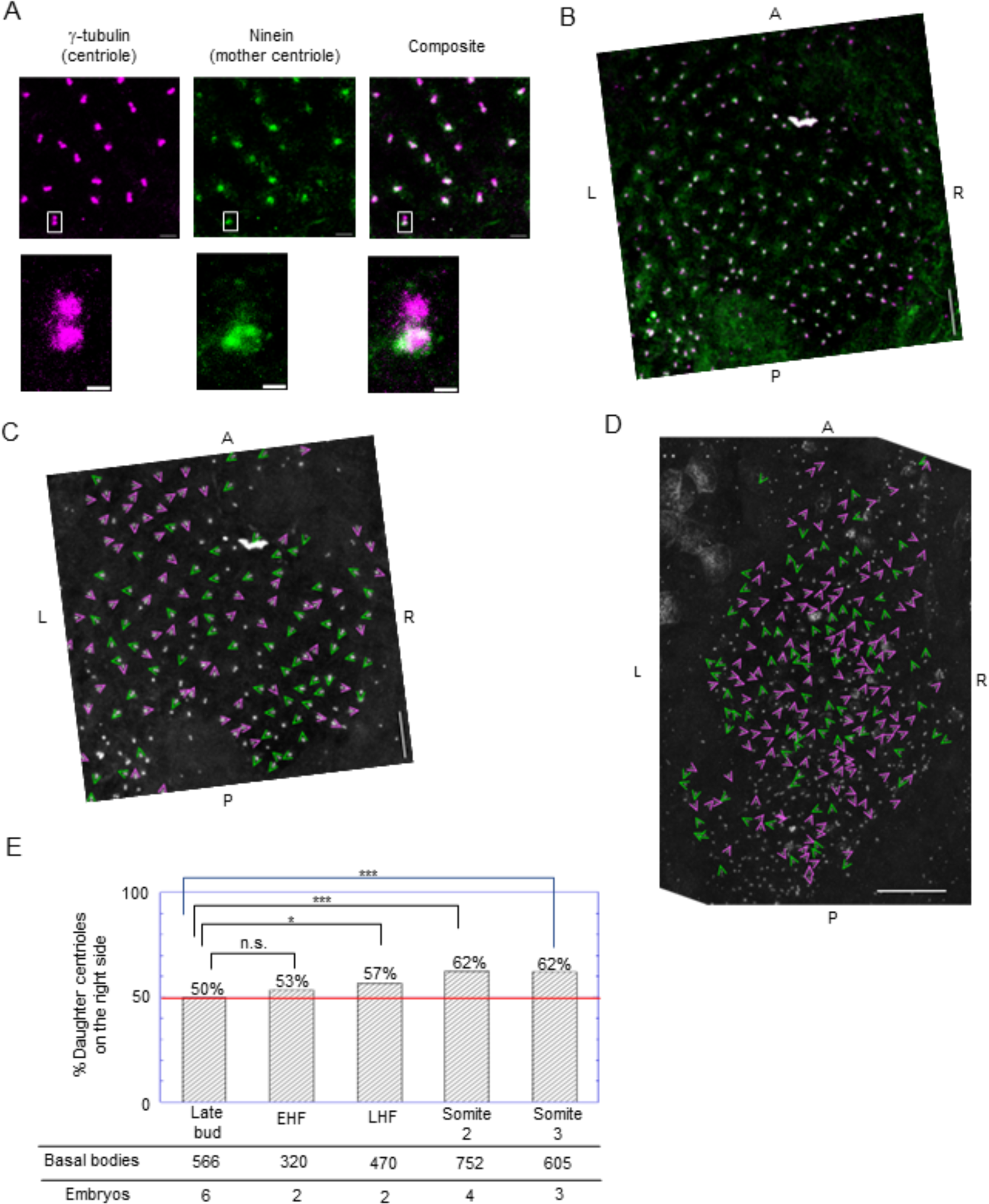
Daughter centrioles become biased to the right side of the mother centrioles in the node in a stage-dependent manner. A. Ninein localizes predominantly to one of the pairs of centrioles in the node of the mouse. XY-projected images (maximum intensity projection) of immunostaining for γ-tubulin (magenta) and ninein (green) in the node are shown. Bars, 2 μm for upper three panels and 500 nm for lower three panels. B. An example (xy-projected image) of the result of immunostaining for γ-tubulin (magenta) and ninein (green) in the node of a late-bud embryo. A, anterior; P, posterior; L, left; R, right. Bar, 10 μm. C. The result of the analysis of the relative positions of the daughter centrioles to the mother centrioles in a late bud embryo shown in B. The directions of the arrowheads indicate the directions of the daughter centrioles from the mother centrioles for each pair of centrioles. The grayscale shows the γ-tubulin signal (shown in magenta in B). In this embryo (Embryo number 1217-#6 in Table 1), the numbers of the daughter centrioles at the right side of the mother centrioles (indicated by magenta arrowheads) and the daughter centrioles at the left side of the mother centrioles (indicated by green arrowheads) were 68 and 66, respectively. Bar, 10 μm. D. The directions of the daughter centrioles from the mother centrioles in the node of a somite-2 stage embryo. In this embryo (#3 in Table 2), the number of daughter centrioles at the right side (magenta arrowheads) and the left side (green arrowheads) were 140 and 81, respectively. Bar, 20 μm. E. The percentage of the daughter centrioles at the right side of the mother centrioles increased in a stage-dependent manner. EHF, early headfold; LHF, late headfold. *, *P*<0.05; ***, *P*<10^-4^; n.s., *P*>0.05; chi-square test. See also Tables 1 and 2.

**Table 1.** No significant L/R asymmetry was detected in the positions of the mother/daughter centrioles in earlier stage embryos.

| Stage | Embryo number | Daughter centrioles on |  | *P* value<br>(binomial test) |  |
| --- | --- | --- | --- | --- | --- |
|  |  | Left | Right |  |  |
| Late bud | 1217-#6 | 66 | 68 | 0.93 | n. s. |
|  | 1222-#4 | 36 | 25 | 0.2 | n. s. |
|  | 0715-#4 | 27 | 23 | 0.67 | n. s. |
|  | 0715-#5 | 6 | 8 | 0.79 | n. s. |
|  | 0502-#1 | 73 | 79 | 0.68 | n. s. |
|  | 0509-#2 | 76 | 79 | 0.87 | n. s. |
| Early headfold | 0715-#8 | 43 | 64 | 0.052 | n. s. |
|  | 0521-#4 | 106 | 107 | 1 | n. s. |

For each embryo, the numbers of daughter centrioles on the left side and the right side of the mother centrioles in the node are shown with the *P* values of the binomial test. n.s., $P > 0.05$ .

**Table 2.**
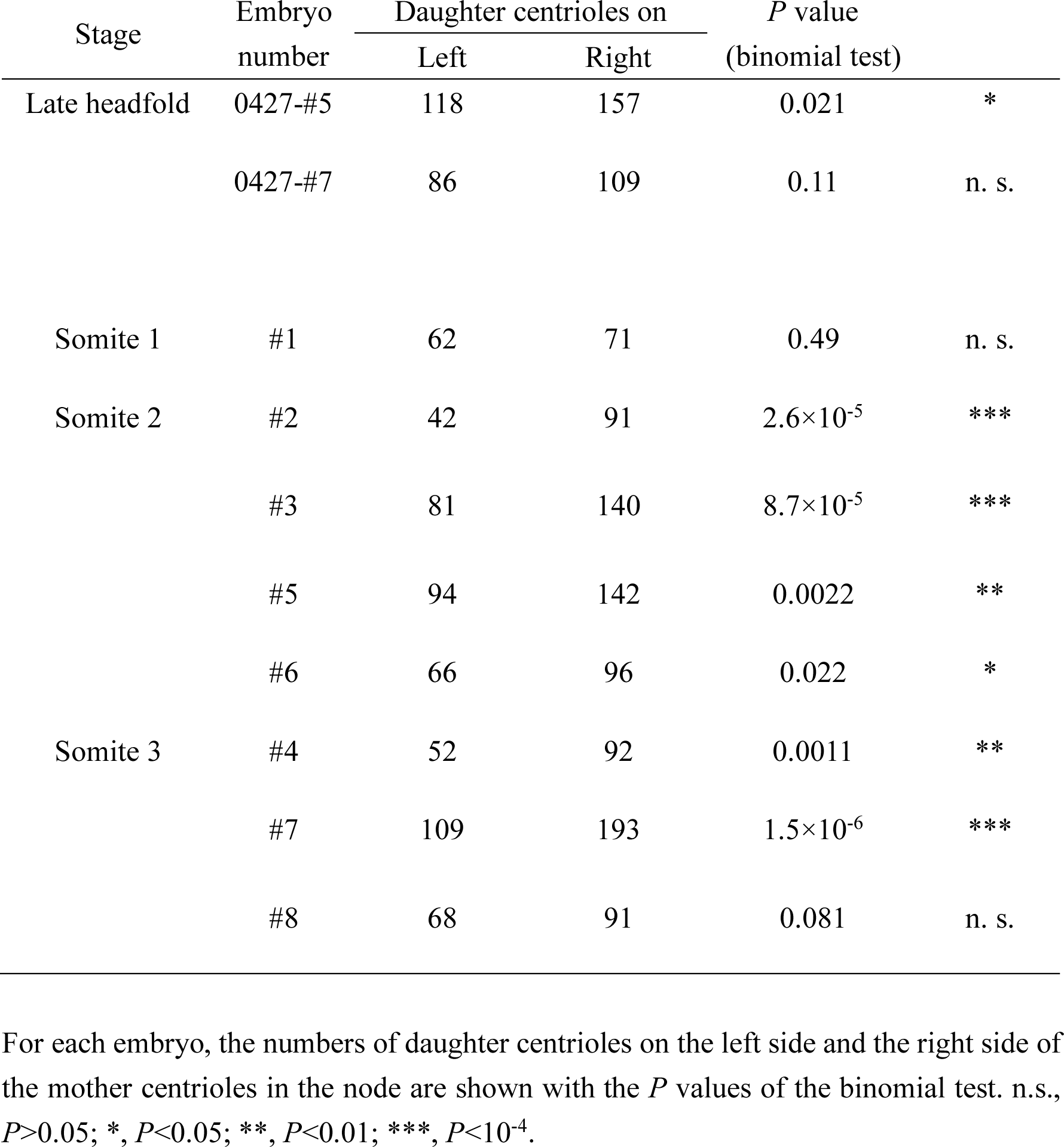
Daughter centrioles were biased to the right side of the mother centrioles in later stage embryos.

### Formation of the L/R asymmetry in the basal bodies in the node requires cilia motility

Since we found that this L/R asymmetry in the basal bodies (i.e., the positions of the daughter centrioles being biased to the right side of the mother centrioles in the node) is formed during developmental stages when node cilia start moving, we then tested whether cilia motility is required for the formation of the L/R asymmetry in the basal bodies. We investigated the positions of the mother and daughter centrioles in the node of *iv/iv* mutant mouse embryos. *iv* (*inversus viscerum*) is a mutation in *left/right-dynein* (*lrd*), which encodes the heavy chain of an axonemal dynein that powers the motility of the node cilia (Supp et al., 1997). The homozygous *iv/iv* mutants lack motility of the node cilia and thus lack fluid flow in the node (Okada et al., 1999). We found that in somite-2 (and somite-3) *iv/iv* embryos, no significant L/R bias was detected in the relative positions of the daughter centrioles to the mother centrioles (Table 3, Fig. 2A). The percentage of the daughter centrioles on the right side in *iv/iv* somite-2 embryos (49%) was comparable to that in wild type late bud embryos (50%), and significantly lower (*P*<10^-4^, chi-square test) than that in wild type somite-2 embryos (62%) (Fig. 2B). These results suggest that formation of the L/R asymmetry in the basal bodies in the node requires cilia motility.

**Fig. 2.**
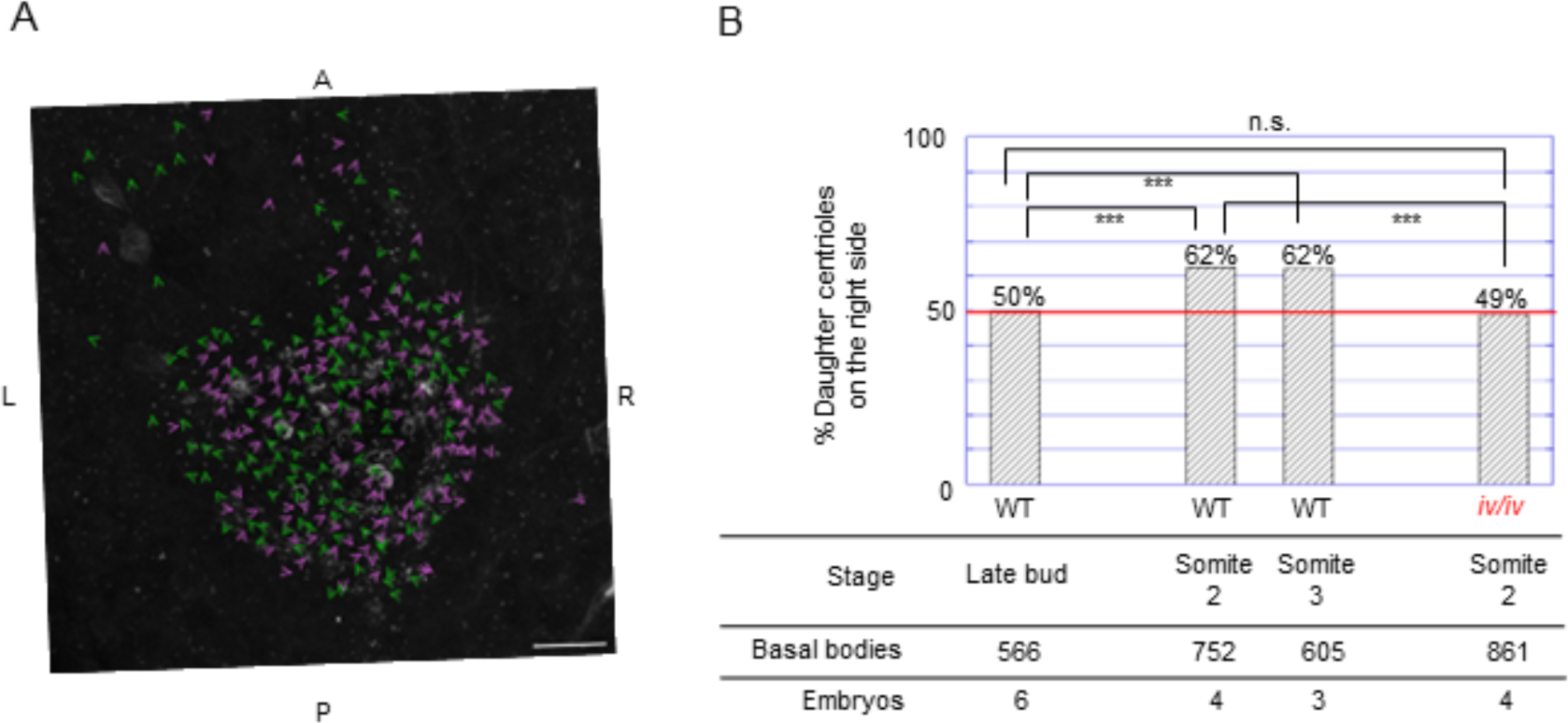
Formation of L/R asymmetry in the basal bodies of the node cilia requires cilia motility. A. The directions of the daughter centrioles from the mother centrioles in the node of an *iv/iv* somite-2 mouse embryo. The arrowheads showing the directions of the daughter centrioles are superimposed on the grayscale image showing γ-tubulin signals. In this *iv/iv* embryo (*iv*-#6 in Table 3), in which node cilia are immotile, the numbers of the daughter centrioles at the right side of the mother centrioles (magenta arrowheads) and the left side of the mother centrioles (green arrowheads) were 133 and 130, respectively. Bar, 20 μm. B. L/R asymmetry of the basal bodies was not detected in *iv/iv* mutant somite-2 mice. ***, *P*<10^-4^; n.s., *P*>0.05; chi-square test. See also Table 3.

**Table 3.** No significant L/R asymmetry was detected in the positions of the mother/daughter centrioles in *iv/iv* embryos.

| Stage | Embryo number | Daughter centrioles on |  | *P* value<br>(binomial test) |  |
| --- | --- | --- | --- | --- | --- |
|  |  | Left | Right |  |  |
| Somite 2 | iv-#3 | 87 | 86 | 1 | n. s. |
|  | iv-#4 | 111 | 87 | 0.10 | n. s. |
|  | iv-#5 | 112 | 115 | 0.89 | n. s. |
|  | iv-#6 | 130 | 133 | 0.90 | n. s. |
| Somite 3 | iv-#8 | 89 | 95 | 0.71 | n. s. |

For each *iv/iv* embryo, the numbers of daughter centrioles on the left side and the right side of the mother centrioles in the node are shown with the *P* values of the binomial test. n.s., $P > 0.05$ .

### Daughter centrioles become biased to the anterior side of the mother centrioles in a cilia motility-independent manner

In the above discussions, we divided the data of the relative positions of the daughter centrioles to two categories: the daughter centrioles at the left side of the mother centrioles and those at the right side of the mother centrioles. To obtain insights into the mechanism of producing the L/R asymmetry in the relative positions of the daughter centrioles, we also focused on the relative positions of the daughter centrioles in the anterior-posterior axis. We found that in wild type somite-2 embryos, the positions of the daughter centrioles tended to be at the anterior side of the mother centrioles (Fig. 3B,E), in addition to the bias toward the right side already shown in a previous section (Fig 3B,E, Fig. 1D,E, Table 2). We then divided these data of the relative positions of the daughter centrioles to two categories: the daughter centrioles at the anterior side of the mother centrioles and those at the posterior side of the mother centrioles. We found that in each of the 4 wild type somite-2 embryos investigated, the daughter centrioles were biased to the anterior side of the mother centrioles significantly (*P*<10^-6^, binomial test; Table 4). Such a clear bias toward the anterior side was not observed in wild type late bud embryos (Table 4, Fig. 3A,D). Furthermore, we found that in *iv/iv* mutants, in which L/R asymmetry of the basal bodies was absent in somite-2 embryos (Table 3, Fig. 2), the bias toward the anterior side was present in somite-2 embryos (Table 4, Fig. 3C,F). The percentage of the daughter centrioles at the anterior side of the mother centrioles increased significantly from late bud to somite 2 in wild type (*P*=5.2×10^-31^, chi-square test), but it was comparable between wild type and *iv/iv* mutants in the somite-2 stage (*P*>0.13, chi-square test; Fig. 3G). These results suggest that the positions of the daughter centrioles in the node become biased to the anterior side of the mother centrioles in a stage-dependent manner, and the formation of the bias toward the anterior side is independent from the cilia motility.

**Fig. 3.**
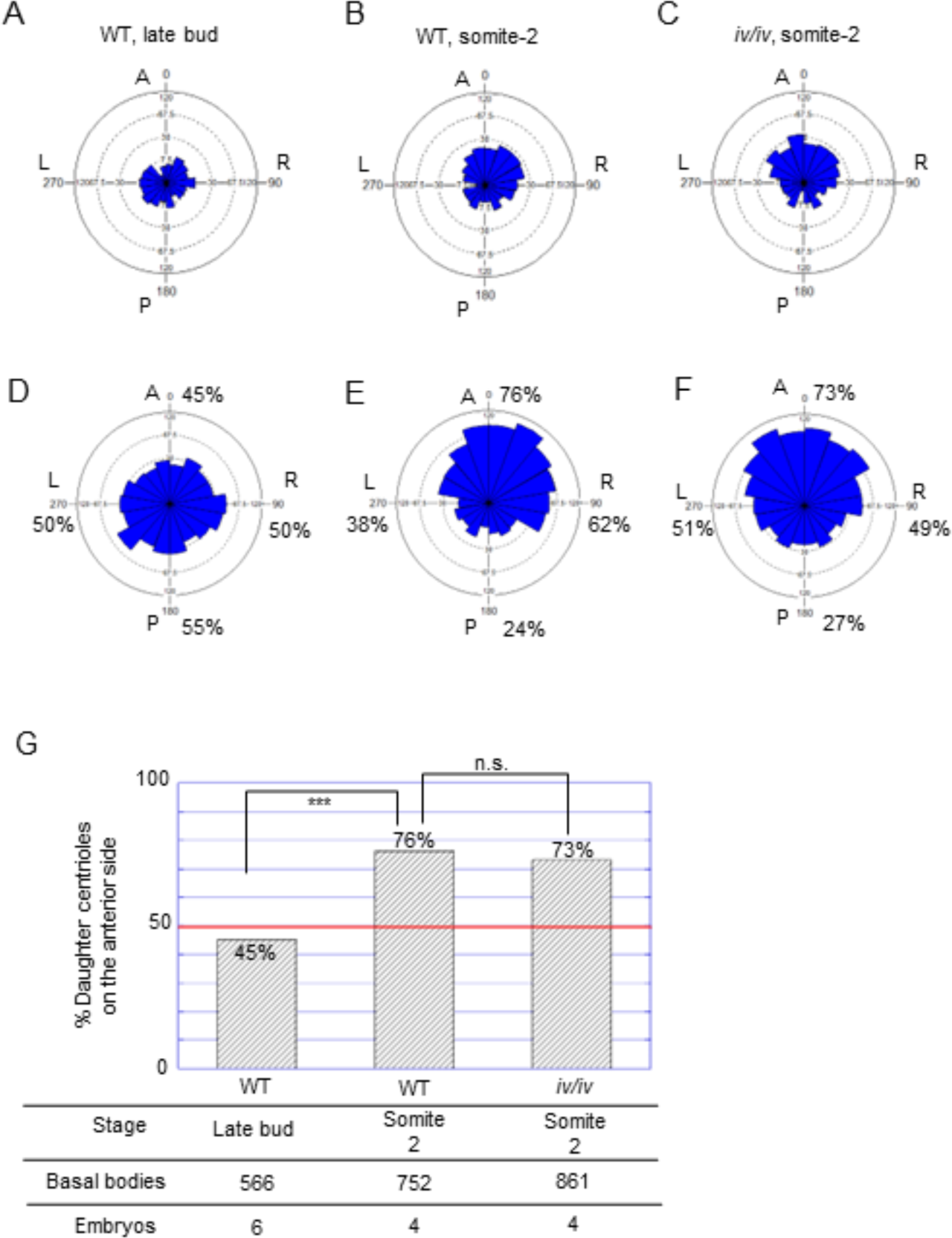
Daughter centrioles become biased to the anterior side of the mother centrioles in a stage-dependent and cilia motility-independent manner. A–C. The rose histograms summarizing the relative positions of the daughter centrioles to the mother centrioles in the node of an embryo from three categories. A, B and C summarize the data from a wild type late bud embryo shown in Fig. 1C, a wild type somite-2 embryo shown in Fig. 1D and an *iv/iv* somite-2 embryo shown in Fig. 2A, respectively. The areas of the wedges indicate the frequencies. D–F. The histograms summarizing the relative positions of the daughter centrioles to the mother centrioles in the node in 6 wild type late bud embryos (D), 4 wild type somite-2 embryos (E) and 4 *iv/iv* somite-2 embryos (F). G. The percentage of the daughter centrioles at the anterior side of the mother centrioles increased from late bud to somite-2 stage, but it was comparable between wild type and *iv/iv* in somite-2 stage. ***, *P*=5.2×10^-31^ <10^-4^; n.s., *P*>0.13 >0.05; chi-square test.

**Table 4.**
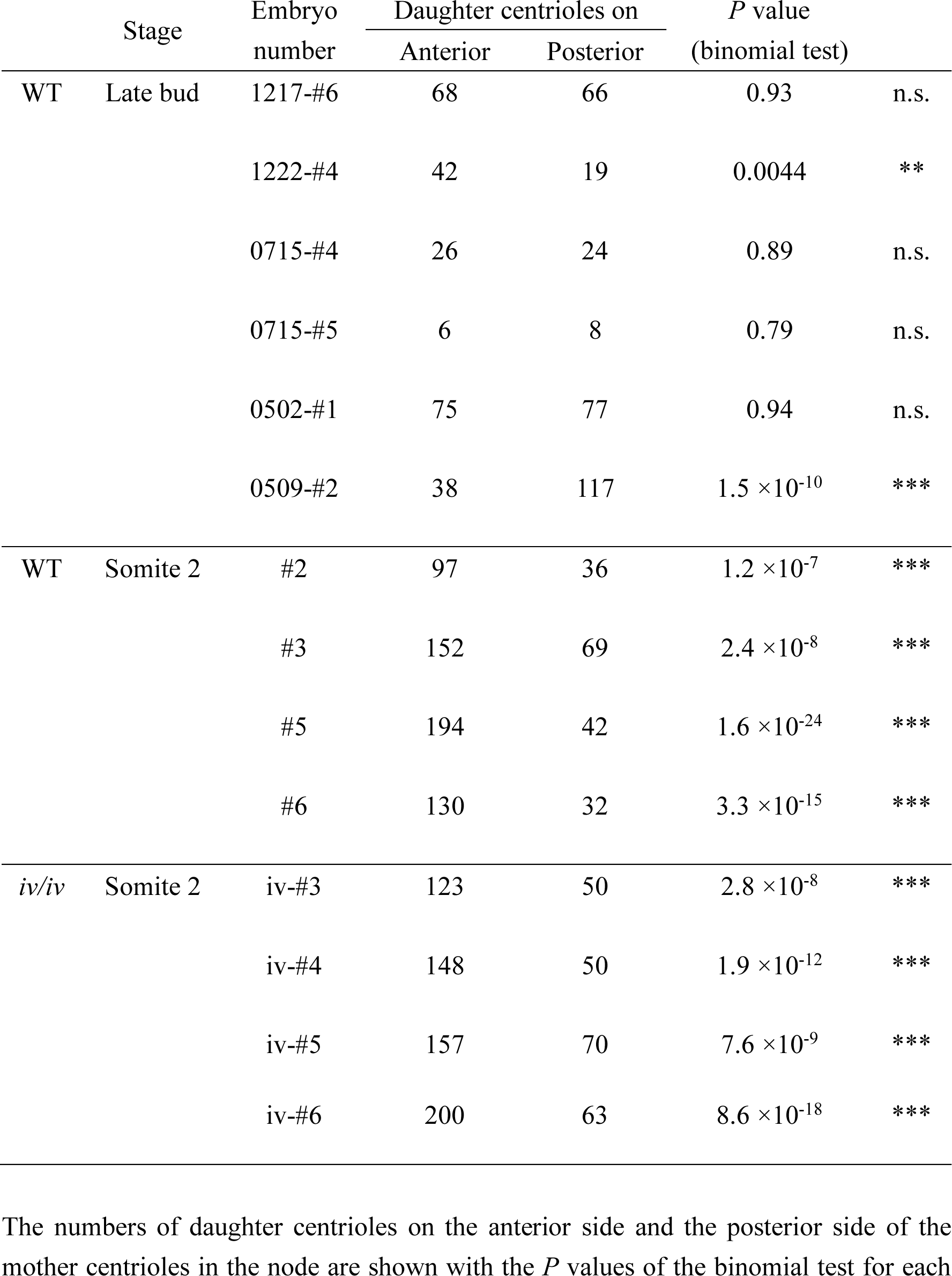
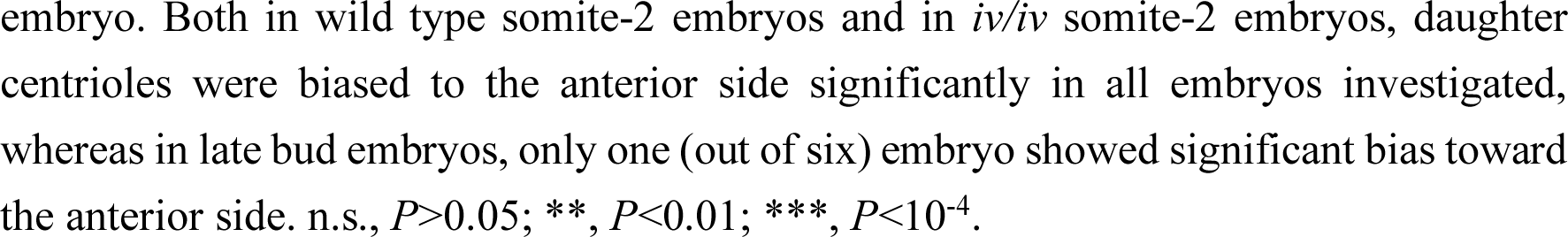
Daughter centrioles were biased to the anterior side of the mother centrioles in wild type somite-2 embryos and also in *iv/iv* somite-2 embryos.

| Stage |  | Embryo number | Daughter centrioles on |  | *P* value<br>(binomial test) |  |
| --- | --- | --- | --- | --- | --- | --- |
|  |  |  | Anterior | Posterior |  |  |
| WT | Late bud | 1217-#6 | 68 | 66 | 0.93 | n.s. |
|  |  | 1222-#4 | 42 | 19 | 0.0044 | ** |
|  |  | 0715-#4 | 26 | 24 | 0.89 | n.s. |
|  |  | 0715-#5 | 6 | 8 | 0.79 | n.s. |
|  |  | 0502-#1 | 75 | 77 | 0.94 | n.s. |
| | | 0509-#2 | 38 | 117 | $1.5 \times 10^{-10}$ | *** |
| WT | Somite 2 | #2 | 97 | 36 | $1.2 \times 10^{-7}$ | *** |
| | | #3 | 152 | 69 | $2.4 \times 10^{-8}$ | *** |
| | | #5 | 194 | 42 | $1.6 \times 10^{-24}$ | *** |
| | | #6 | 130 | 32 | $3.3 \times 10^{-15}$ | *** |
| *iv/iv* | Somite 2 | *iv*-#3 | 123 | 50 | $2.8 \times 10^{-8}$ | *** |
| | | *iv*-#4 | 148 | 50 | $1.9 \times 10^{-12}$ | *** |
| | | *iv*-#5 | 157 | 70 | $7.6 \times 10^{-9}$ | *** |
| | | *iv*-#6 | 200 | 63 | $8.6 \times 10^{-18}$ | *** |

The numbers of daughter centrioles on the anterior side and the posterior side of the mother centrioles in the node are shown with the *P* values of the binomial test for each
embryo. Both in wild type somite-2 embryos and in *iv/iv* somite-2 embryos, daughter centrioles were biased to the anterior side significantly in all embryos investigated, whereas in late bud embryos, only one (out of six) embryo showed significant bias toward the anterior side. n.s., $P>0.05$ ; \*\*, $P<0.01$ ; \*\*\*, $P<10^{-4}$ .

It has been reported that basal bodies of the node cilia translocate from the center of the cells toward the posterior side of the cells during stages including the period from late bud to somite-2 (Hashimoto et al., 2010). This posterior translocation is based on anterior-posterior (A-P) polarization of the node cells mediated by noncanonical Wnt signaling pathway including Wnt5, Rac1, Dvl and PCP (planar cell polarity) core components including Prickle1 and Prickle2 (Hashimoto et al., 2010; Minegishi et al., 2017). The present result that the daughter centrioles become biased to the anterior side of the mother centrioles during this period suggests that during this translocation of the basal bodies, the mother centrioles lead (move first toward the posterior side of the cells) and the daughter centrioles follow. This implies that the relative positions of the daughter centrioles could be affected by A-P polarization of the node cells and by posterior translocation of the basal bodies independently from the cilia motility, and raises up a possibility that formation of L/R asymmetry of the basal bodies could also be affected by the preexisting A-P axis information established independently from the cilia motility.

### L/R asymmetry of the basal bodies was not reversed by artificial counter-flow

We found that L/R asymmetry of the basal bodies was not formed in *iv/iv* somite-2 mutants (Table3, Fig. 2), suggesting that the formation of L/R asymmetry of the basal bodies requires cilia motility. However, since *iv/iv* embryos lack cilia motility and also lack leftward nodal flow resulting from the motility of the node cilia, we did not know whether the motility of the cilia themselves are required, or the resultant leftward fluid flow is required. To distinguish between these possibilities, we tested whether the application of artificial counter-flow, which reverses the direction of the fluid flow in the node, can reverse the L/R asymmetry of the basal bodies. Nonaka et al. showed that by culturing mouse embryos in a flow chamber with flow of the culturing medium toward the opposite direction to the endogenous leftward nodal flow, the direction of the fluid flow in the node can be reversed, and the L/R asymmetry in the heart loops and the embryonic turning in later stages was also reversed (Nonaka et al., 2002). First, we reexamined whether the artificial counter-flow can reverse the chirality of the heart loops. Pre-somite embryos were cultured with artificial counter-flow overnight, and further cultured by conventional rotation culture method for 32–33 h. Out of six embryos investigated, the hearts of all of them showed a chirality called “L-loop” (Fig. 4B,C), which is the opposite to that of normal development (“D-loop”, Franco et al., 2001). Although in 4 out of 6 embryos showed no embryonic turning and the tails were at the back side of the embryos, in 2 of them in which embryonic turning has occurred, tails were at the left side of the embryos (Fig. 4A,B), which is the opposite to the normal development. Thus, we confirmed that artificial counter-flow can reverse the L/R asymmetric morphogenesis including the chirality of the heart loops.

**Fig. 4.**
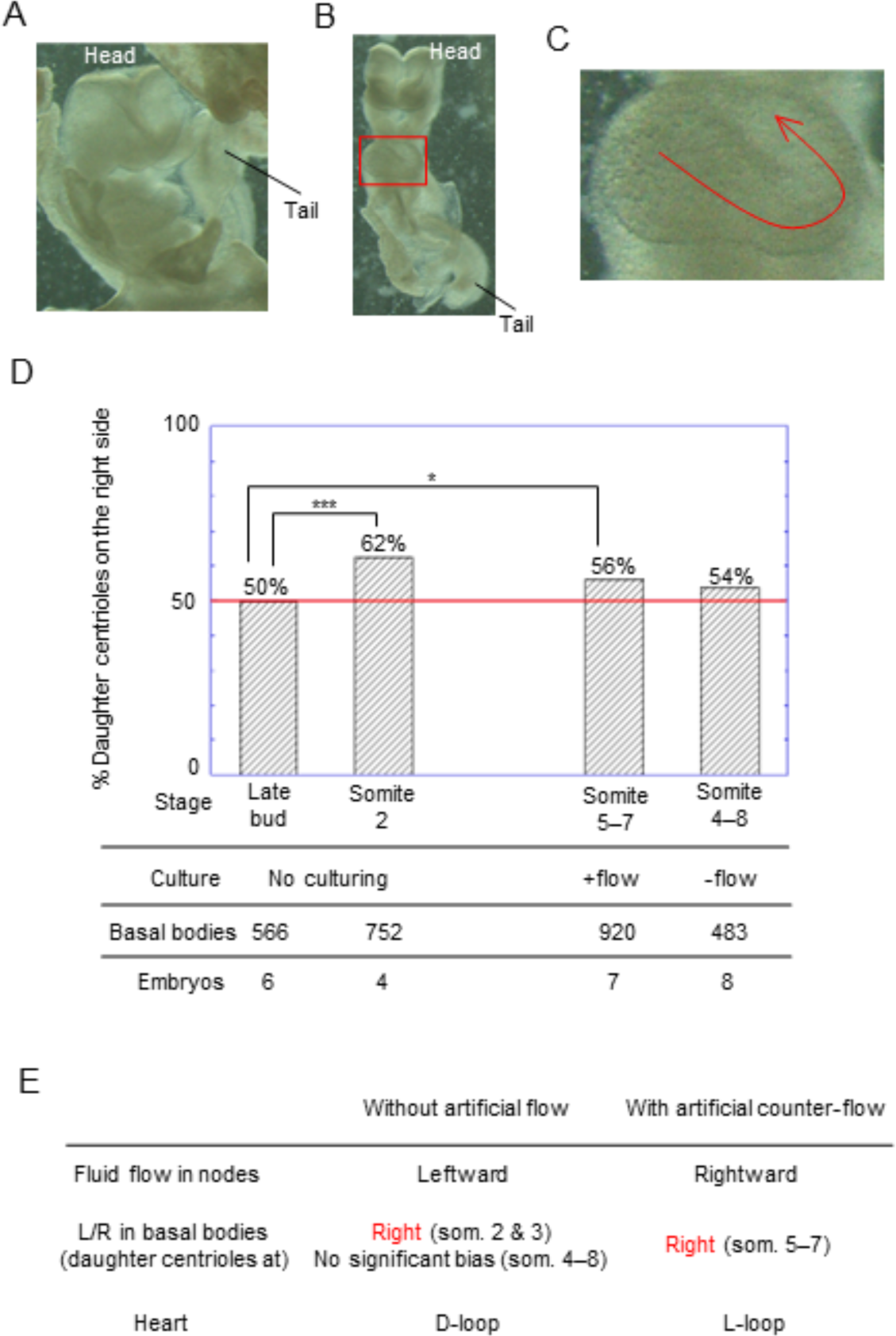
L/R asymmetry of the basal bodies was not reversed by artificial counter-flow. A–B. Stereomicroscopic images of the embryos cultured with an artificial counter-flow overnight followed by conventional rotation culture for 32–33 h. Axial turning has begun and the tails are at the left side of the embryos, which is the opposite side to the normal development. C. A magnified image of the heart (the rectangle shown in B). The cardiac tube loops toward the left side of the embryo (L-loop), which is the opposite to the chirality in the normal development (D-loop; Franco et al., 2001). D. The percentage of the daughter centrioles at the right side of the mother centrioles in the node was significantly higher in the somite 5–7 embryos which were cultured with an artificial counter-flow overnight than in late bud embryos. “No culturing”, embryos were dissected from pregnant mice and fixed for immunostaining without culturing; “+flow”, pre-somite stage embryos were cultured with an artificial counter-flow overnight; “-flow”, pre-somite stage embryos were cultured in a conventional rotation culture overnight without applying artificial flow. *, *P*<0.05; ***, *P*<10^-4^; chi-square test. See also Table 4. E. Summary of the result of the artificial flow experiment.

Then, we tested whether the artificial counter-flow of the same condition can reverse the L/R asymmetry of the basal bodies. Interestingly, we found that out of 7 embryos immunostained after application of artificial counter-flow overnight, none showed left-sided bias in the positions of the daughter centrioles (Table 5). On the contrary, in 3 out of 7 embryos, the positions of the daughter centrioles were significantly biased to the right side of the mother centrioles (*P*<0.05, binomial test, Table 5). Also, the total number of the daughter centrioles on the right side of the mother centrioles in the 7 embryos investigated was significantly more than that on the left side of the mother centrioles (*P*=1.9×10^-4^, binomial test, Table 5). The percentage of the daughter centrioles on the right side of the mother centrioles in the node of the embryos cultured with artificial counter-flow overnight (56%) was significantly higher than that of late bud embryos (50%) (*P*<0.05, chi-square test, Fig. 4D). These results suggest artificial counter-flow, which reversed the L/R asymmetry of the heart loops, did not reverse the L/R asymmetry of the basal bodies, and the daughter centrioles still tended to be at the right side of the mother centrioles (Fig. 4E). Combined with the result of the analysis of *iv/iv* mutants (Table 3, Fig. 2), this result suggests that the formation of the L/R asymmetry of the basal bodies requires cilia motility itself, and that it is independent from the direction of the fluid flow in the node. It is also suggested that the L/R asymmetry of the basal bodies is not the determinant of the L/R asymmetry of the heart loops (Fig. 4E).

**Table 5.** Daughter centrioles tended to be at the right side of the mother centrioles in the node even after application of artificial counter-flow.

| Embryo number | Somites | Daughter centrioles on |  | *P* value<br>(binomial test) |  |
| --- | --- | --- | --- | --- | --- |
|  |  | Left | Right |  |  |
| 0414-#4 | 5 | 76 | 109 | 0.018 | * |
| 0414-#6 | 5 | 98 | 101 | 0.89 | n.s. |
| 0414-#1 | 6 | 37 | 64 | 0.0093 | ** |
| 0414-#3 | 6 | 20 | 24 | 0.65 | n.s. |
| 0414-#7 | 6 | 38 | 60 | 0.033 | * |
| 0414-#5 | 6 | 46 | 48 | 0.92 | n.s. |
| 0414-#2 | 7 | 88 | 111 | 0.12 | n.s. |
| Total | | 403 | 517 | $1.9 \times 10^{-4}$ | ** |

For each embryo to which artificial counter-flow was applied overnight, the numbers of somites after the application of artificial flow and the numbers of daughter centrioles on the left side and the right side of the mother centrioles in the node are shown with the *P* values of the binomial test. The total numbers of centrioles are also shown with the *P* value of the binomial test. n.s., $P > 0.05$ ; \*, $P < 0.05$ ; \*\*, $P < 0.01$ .

## Discussion

In this study, we found that the positions of the daughter centrioles beside the mother centrioles at the base of the node cilia become biased to the right side of the mother centrioles in a stage-dependent manner, and that formation of this L/R asymmetry in the basal bodies is dependent on cilia motility but does not depend on the direction of the fluid flow in the node. It is well established that the node cilia produce L/R asymmetry by producing leftward fluid flow (Okada et al., 1999; Okada et al., 2005), and the leftward fluid flow is responsible for the L/R asymmetric gene expressions and morphogenesis (Lowe et al., 1996; Nonaka et al., 2002). The present results suggest that the node cilia may have another mechanism of producing L/R asymmetry, in the basal bodies inside the cells independently from the fluid flow in the node.

The mechanism of how the daughter centrioles become biased to the right side of the mother centrioles via cilia motility is unknown, but we speculate torque imposed on the basal bodies as a result of the rotational movement of the cilia themselves may promote rotation of the centrioles at the base to form right-sided arrangement of the daughter centrioles. As a support to this speculation, Ferreira et al. estimated the torque caused by active motion of the zebrafish Kupffer’s vesicle cilia (the time-averaged meridional component of the torque having a magnitude of about 10^-17^ Nm) and discussed that cells sensing the direction of the torque at the base of the cilium generated by the motion of its own cilium is a plausible mechanism for a L/R asymmetric response in the Kupffer’s vesicle in zebrafish (Ferreira et al., 2017).

In contrast to the previously known mechanism of producing L/R asymmetry via the fluid flow in the node, the novel mechanism found in this study is likely to be independent from the L/R asymmetry of the visceral organs in mice, because embryos cultured with artificial counter-flow overnight showed reversed chirality of the heart loops, but did not show reversed arrangements in the relative positions of the daughter centrioles (Fig. 4). Thus, at this moment, it is unknown whether this L/R asymmetry in the basal bodies has any biological significance. However, we consider this novel mechanism of producing L/R asymmetry via cilia motility could be a candidate for a symmetry breaking mechanism in two cases in which initial L/R production mechanism has been mysterious.

One is determination of L/R asymmetry in the visceral organs in other animals such as chicks and pigs. In rabbits, frogs, and fish, rotational ciliary movement and directional fluid flow in the left-right organizer have been observed (Okada et al., 2005; Schweickert et al., 2007), disruption of fluid flow by methyl cellulose results in defects in laterality in the visceral organs in frogs (Schweickert et al., 2007), and inhibition of cilia motility by knockdown of ciliary dynein gene *lrdr1* results in perturbation of L/R asymmetric gene expressions (Essner et al., 2005). In contrast, in chicks, although several kinds of L/R asymmetry have been observed around the primitive streak and Hensen’s node (left-right organizer) such as asymmetric membrane potential (Levin et al., 2002), asymmetric concentration of extracellular Ca^2+^ (Raya et al., 2004), and rotational movement of cells around Hensen’s node (Gros et al., 2009), the initial symmetry breaking event at the upstream of those asymmetric phenomena has been unknown. In chicks and pigs, unlike in mice, the ventral side of Hensen’s node is covered by endoderm cells and appears to have no space for fluid flow (Essner et al., 2002; Stephen et al., 2014; Gros et al., 2009), so fluid flow-dependent symmetry breaking may not regulate laterality in chicks and pigs. The epiblast (dorsal side) nodal pit cells and mesoderm cells at Hensen’s node in chicks sparsely contain short monocilia (Männer, 2001; Stephen et al., 2014). These cilia in the Hensen’s node have been suggested to be probably immotile based on low expression of *FOXJ1*, a master regulator of motile cilia (Stephen et al., 2014), but to our knowledge, the presence/absence of motility in these cilia have not been observed directly, and at least the ciliary dynein gene *Lrdr* is expressed within Hensen’s node in the chick (Essner et al., 2002). Also, at the margin of the epiblast nodal pit and in the more peripheral regions, cilia are more frequently observed (Männer, 2001; Stephen et al., 2014). These cilia have been previously overlooked because they are not likely to produce fluid flow within the pit, but the present study raises up a possibility that these cilia might also have a capacity to develop L/R asymmetry in the basal bodies, if they are motile. It is an open question whether these cilia are motile or not, whether the positions of the mother centrioles and the daughter centrioles at the base of cilia have L/R asymmetry, and if so, whether it is relevant with the previously reported L/R asymmetric phenomena and the laterality of the organs.

Another is the determination of L/R asymmetry in the hippocampus in the brains in the mouse. It has been reported that distribution of *N*-methyl-D-aspartate (NMDA) receptor GluRχ2 subunits in synapses in the adult mouse hippocampus has L/R asymmetry (Kawakami et al., 2003). This L/R asymmetry in the brain is lost in *iv/iv* mice, suggesting that formation of this L/R asymmetry requires a function of the same ciliary dynein gene (*lrd)* as is required for L/R determination of the visceral organs (Kawakami et al., 2008). However, while visceral organs show randomized laterality in *iv/iv* mice, allocation of GluRχ2 subunits in the hippocampus always showed right isomerism (i.e., both left and right sides showed the right side-type phenotype) regardless of the laterality of the visceral organs (*situs solitus* or *situs inversus*) in *iv/iv* mice (Kawakami et al., 2008). This suggests that distinct mechanism from visceral organs generates L/R asymmetry in the brains, but the mechanism is unknown. Now, we propose that the L/R asymmetry in the basal bodies in the node could be a candidate of a determinant of the L/R asymmetry in the brains, because they have three characteristic features in common: existence of L/R asymmetry in wild type; lack of L/R asymmetry in *iv/iv* mice; and being independent from the L/R asymmetry of the visceral organs. *Lrd* gene, which is mutated in *iv* mice, is not only expressed in the node but also in gut, limbs and the floorplate in the neural tubes in later stages (Supp et al., 1999; Cruz et al., 2010), so we cannot exclude a possibility of involvement of dynein in other tissues than the node in determining L/R asymmetry in the hippocampus. However, the functions of the dynein heavy chain encoded by *lrd* in other tissues are unknown, and it is also possible that *lrd* expressed in the node and powering the motility of the node cilia could function in determining the L/R asymmetry in the hippocampus of the mouse.

In any cases, the pathway or the mechanism connecting the L/R asymmetry in the basal bodies in the node and a possible downstream L/R asymmetric phenomenon is totally unknown, but the present study provides a new possibility that the node cilia may initiate L/R asymmetry in a “non-canonical” mechanism independent of the fluid flow, and may provide a clue to better understand the diversity in the mechanisms of creating L/R asymmetry in vertebrates.

## Materials and Methods

### Mouse embryos

ICR mice purchased from Japan SLC Inc. were mated to obtain wild type embryos. *iv/iv* mice, described previously (Supp et al., 1997) were mated to obtain *iv/iv* embryos. Embryos were dissected from pregnant mice at embryonic day 7.7–7.8 (wild type) or 8.0 (*iv/iv*) in Dulbecco’s Modified Eagle Medium (DMEM, 12320-032, Gibco) supplemented with 10% fetal bovine serum (FBS, 26140-079, Gibco). Embryos were staged on the basis of the morphology (Downs and Davies, 1993).

### Immunofluorescence

Embryos were fixed with methanol (21923-25, nacalai) on ice for 10 min. After washing with PBS (phosphate-buffered saline) for three times, embryos were soaked in 1% BSA (bovine serum albumin, A9647, SIGMA) in PBS in room temperature for 30 min for blocking. Then embryos were incubated with primary antibodies for 1 h in room temperature or overnight in 4℃. Primary antibodies used were rabbit anti-ODF2 polyclonal antibody (ab43840, abcam), mouse anti-centriolin monoclonal antibody (sc-365521, Santa Cruz), rabbit anti-ninein polyclonal antibody (PA5-96974, Invitrogen) and mouse anti-γ-tubulin monoclonal antibody (T5326, Sigma-Aldrich). Then embryos were washed with PBS for three times, and incubated with the secondary antibodies. The secondary antibodies used for double staining were either the combination of Alexa Fluor 594-conjugated anti-rabbit IgG antibody (ab150080, abcam) and Alexa Fluor 647-conjugated anti-mouse IgG antibody (a21237, Invitrogen), or the combination of Oregon green 488-conjugated anti-rabbit IgG antibody (O-11038, Invitrogen) and Cyanine 3-conjugated anti-mouse IgG antibody (M30010, Invitrogen). Embryos were washed with PBS for three times and then PBS was replaced with 10%, 25%, 50% and 97% TDE (2,2’-thiodiethanol, 166782, Sigma-Aldrich), sequentially. 10%, 25% and 50% TDE were obtained by diluting 97% (v/v) TDE with 4% (w/w) propyl gallate (162-06832, Wako) by PBS to the ratio of 1:10, 1:4 and 1:2, respectively. The procedures described in this paragraph were performed in a 24-well microplate (1820-024, IWAKI).

Preparation of the sample for imaging was performed as described previously (Fig. 2 in Katoh et al., 2023). The stained embryos were cut using two needles and the fragment of the embryo containing the node was carefully placed in the hole of a silicone rubber spacer (0.25 mm thick) fitted on a glass slide (S1111, Matsunami). The embryo and the spacer were covered with a coverslip (No. 1SHT, Matsunami) and sealed with nail polish.

Images were acquired using TCS SP8 STED 3X FALCON (Leica) with HC PL APO 100x/1.40 oil-immersion STED objective or TCS SP8 MP (inverted, Leica) with HC PL APO 63x/1.40 oil-immersion objective. When TCS SP8 STED 3X FALCON was used, images, including those in Fig. 1A and Fig. S1, were acquired using STED (stimulated emission depletion) microscopy (Klar et al., 2000). 775 nm STED laser was used for depletion of Alexa Fluor 594 and Alexa Fluor 647, and 660 nm STED laser was used for Oregon green 488 and Cyanine 3. Please note that super-resolution of STED microscopy was not necessary for the analysis of the relative positions of the centrioles in this study. The resolution of the confocal microscope by TCS SP8 MP was sufficient for discriminating the positions of the mother and daughter centrioles, which are hundreds of nanometers (or more) apart from each other.

### Analysis of the positions of the mother and daughter centrioles

XY plane-projected images of the confocal z-stack images with maximum intensity projection were created with the image analysis software ImageJ (Abramoff et al., 2004). The coordinates of the mother and daughter centrioles in the XY-projected images were obtained by manually clicking on the centrioles in the images using the multi-point tool of the ImageJ. The A-P (anterior-posterior) axis of the embryos in the images were determined from the morphology of the embryos including the notochord and the node observed in scan-DIC (differential interference contrast) images acquired along with confocal images using TCS SP8. The relative positions of the daughter centrioles from the mother centrioles were calculated from the coordinates, and the arrowheads (as in Fig. 1CD) were drawn on to the images using a macro of ImageJ. Bar graphs were created using KaleidaGraph (Synergy Software) and rose histograms were created with Oriana (Kovach Computing Services).

### Application of artificial counter-flow

The flow culture system was prepared as described previously (Nonaka et al., 2002). The flow chamber made of acrylic with small holes on the bottom to place the embryos was sealed with a gasket made of 1 mm-thick silicone rubber sheet and with a glass slide (S1111, Matsunami), which were held tight with binder clips (19 mm, KOKUYO). The cotton for filter in the flow chamber was obtained from a disposable pippete (7002-002, Iwaki). Artificial flow was induced by a peristaltic pump (MP-1000, Eyela) with silicone tubes (Part No. 125470, Eyela) and 15 ml centrifugation tubes as depulsators.

The embryos between late bud and late headfold were used. They were placed in the holes at the bottom of the flow chamber with an adequate orientation so that artificial flow can be applied to the node toward the right side of the embryos. The embryos were cultured with flow for 16 h. The flow rate of the pump was 2.4×10^2^ ml/h, which is equivalent to the average speed in the flow chamber of 1.4 mm/s. Then the embryos were either fixed for immunofluorescence analysis of the centrioles, or subjected to conventional rotation culture (Nagy et al., 2003) for an additional 32–33 h in a 50 ml centrifugation tube on a rotator (RT-50, Taitec) for observation of the heart loops and the tails. Throughout culture, embryos were maintained at 37℃ under 5.0% CO_2_ in culture medium containing 50% rat serum (THE JACKSON LABORATORY JAPAN) and 50% DMEM without HEPES buffer (11995-065, Gibco).

## Acknowledgement

This work was supported by Optics and Imaging Facility, NIBB Trans-Scale Biology Center.

## References

Abramoff, M. D., Magalhaes, P. J., Ram, S. J. (2004). Image Processing with ImageJ. Biophotonics International 11 (7), 36–42.

Choksi, S. P., Lauter, G., Swoboda, P. and Roy, S. (2014). Switching on cilia: transcriptional networks regulating ciliogenesis. Development 141, 1427–1441.

Cruz, C, Ribes, V., Kutejova, E., Cayuso, J., Lawson, V., Stevens, D. J., Davey, M., Blight, K., Bangs, F., Mynett, A. et al. (2010). Foxj1 regulates floor plate cilia architecture and modifies the response of cells to sonic hedgehog signalling. Development 137, 4271–4282.

Downs, K. M. and Davies, T. (1993). Staging of gastrulating mouse embryos by morphological landmarks in the dissecting microscope. Development 118, 1255–1266.

Essner, J. J., Vogan, K. J., Wagner, M. K., Tabin, C. J., Yost, H. J. and Brueckner, M. (2002). Conserved function for embryonic nodal cilia. Nature 418, 37-38.

Essner, J. J., Amack, J. D., Nyholm, M. K., Harris, E. B. and Yost, H. J. (2005). Kupffer’s vesicle is a ciliated organ of asymmetry in the zebrafish embryo that initiates left-right development of the brain, heart and gut. Development 132, 1247–1260.

Ferreira R. R., Vilfan, A., Jülicher, F., Supatto, W. and Vermot, J. (2017). Physical limits of flow sensing in the left-right organizer. eLIFE 6:e25078.

Fewell, R. M. and Dutcher, S. K. (2020). Basal Feet: Walking to the Discovery of a Novel Hybrid Cilium. Dev. Cell 55, 115–117.

Franco, D., Kelly, R., Moorman, A. F. M., Lamers, W. H., Buckingham, M. and Brown, N. A. (2001). MLC3F Transgene Expression in *iv* Mutant Mice Reveals the Importance of Left-Right Signalling Pathways for the Acquisition of Left and Right Atrial But Not Ventricular Compartment Identity. Dev. Dynam. 221, 206–215.

Gros, J., Feistel, K., Viebahn, C., Blum, M. and Tabin, C. J. (2009). Cell Movements at Hensen’s Node Establish Left/Right Asymmetric Gene Expression in the Chick. Science 324, 941–944.

Hagiwara, H., Ohwada, N., Aoki, T., Suzuki, T. and Takata K. (2008). The primary cilia of secretory cells in the human oviduct mucosa. Med. Mol. Morphol. 41, 193–198.

Hashimoto, M., Shinohara, K., Wang, J., Ikeuchi, S., Yoshiba, S., Meno, C., Nonaka, S., Takada, S., Hatta, K., Wynshaw-Boris, A. and Hamada, H. (2010). Planar polarization of node cells determines the rotational axis of node cilia. Nature Cell Biol. 12, 170–176.

Katoh, T. A., Omori, T., Ishikawa, T., Okada, Y. and Hamada, H. (2023). Biophysical Analysis of Mechanical Signals in Immotile Cilia of Mouse Embryonic Nodes Using Advanced Microscopic Techniques. Bio-protocol 13(14):e4715.

Kawakami, R., Shinohara, Y., Kato, Y., Sugiyama, H., Shigemoto, R. and Ito, I. (2003). Asymmetrical Allocation of NMDA Receptor ε2 Subunits in Hippocampal Circuitry. Science 300, 990–994.

Kawakami, R., Dobi, A., Shigemoto, R. and Ito, I. (2008). Right Isomerism of the Brain in Inversus Viscerum Mutant Mice. PLoS ONE 3(4): e1945.

Klar, T. A., Jakobs, S., Dyba, M., Egner, A. and Hell, S. W. (2000). Fluorescence microscopy with diffraction resolution barrier broken by stimulated emission. P. Natl. Acad. Sci. USA. 97, 8206–8210.

Kobayashi, T. and Dynlacht, B. D. (2011). Regulating the transition from centriole to basal body. J. Cell Biol. 193, 435–444.

Kunimoto, K., Yamazaki, Y., Nishida, T., Shinohara, K., Ishikawa, H., Hasegawa, T., Okanoue, T., Hamada, H., Noda, T., Tamura, A. et al. (2012). Coordinated Ciliary Beating Requires Odf2-Mediated Polarization of Basal Bodies via Basal Feet. Cell 148, 189–200.

Levin, M., Thorlin, T., Robinson, K. R., Nogi, T. and Mercola, M. (2002). Asymmetries in H^+^/K^+^-ATPase and Cell Membrane Potentials Comprise a Very Early Step in Left-Right Patterning. Cell 111, 77–89.

Little, R. B. and Norris, D. P. (2021). Right, left and cilia: How asymmetry is established. Semin. Cell Dev. Biol. 110, 11–18.

Lowe, L. A., Supp, D. M., Sampath, K., Yokoyama, T., Wright, C. V. E., Potter, S. S., Overbeek, P. and Kuehn, M. R. (1996). Conserved left-right asymmetry of nodal expression and alterations in murine *situs inversus*. Nature 381, 158–161.

Männer, J. (2001). Does an equivalent of the “ventral node” exist in chick embryos? A scanning electron microscopic study. Anat. Embryol. 203, 481–490.

Minegishi, K., Hashimoto, M., Ajima, R., Takaoka, K., Shinohara, K., Ikawa, Y., Nishimura, H., McMahon, A. P., Willert, K., Okada, Y. et al. (2017). A Wnt5 Activity Asymmetry and Intercellular Signaling via PCP Proteins Polarize Node Cells for Left-Right Symmetry Breaking. Dev. Cell 40, 439–452.

Minegishi, K., Rothé, B., Komatsu, K. R., Ono, H., Ikawa, Y., Nishimura, H., Katoh, T. A., Kajikawa, E., Sai, X., Miyashita, E. et al. (2021). Fluid flow-induced left-right asymmetric decay of Dand5 mRNA in the mouse embryo requires a Bicc1-Ccr4 RNA degradation complex. Nat. Commun. 12:4071.

Mitchel, B., Jacobs, R., Li, J., Chien, S. and Kintner, C. (2007). A positive feedback mechanism governs the polarity and motion of motile cilia. Nature 447, 97–101.

Nagy, A., Gertsenstein, M., Vintersten, K. and Behringer, R. (2003) Manipulating The Mouse Embryo: A Laboratory Manual Third Edition. Cold Spring Harbor, New York, USA: Cold Spring Harbor Laboratory Press.

Nonaka, S., Tanaka, Y., Okada, Y., Takeda, S., Harada, A., Kanai, Y., Kido, M. and Hirokawa, N. (1998). Randomization of Left–Right Asymmetry due to Loss of Nodal Cilia Generating Leftward Flow of Extraembryonic Fluid in Mice Lacking KIF3B Motor Protein. Cell 95, 829–837.

Nonaka, S., Shiratori, H., Saijoh, Y. and Hamada, H. (2002). Determination of left– right patterning of the mouse embryo by artificial nodal flow. Nature 418, 96–99. et al., 2002

Nonaka, S., Yoshiba, S., Watanabe, D., Ikeuchi, S., Goto, T., Marshall, W. F. and Hamada, H. (2005). De Novo Formation of Left–Right Asymmetry by Posterior Tilt of Nodal Cilia. PLoS Biol. 3(8): e268.

Odate, T., Takeda, S., Narita, K. and Kawahara, T. (2016). 9 + 0 and 9 + 2 cilia are randomly dispersed in the mouse node. Microscopy 65, 119–126.

Okada, Y., Nonaka, S., Tanaka, Y., Saijoh, Y., Hamada, H. and Hirokawa, N. (1999). Abnormal Nodal Flow Precedes Situs Inversus in *iv* and *inv* mice. Mol. Cell 4, 459–468.

Okada, Y., Takeda, S., Tanaka, Y., Izpisúa Belmonte, J.-C. and Hirokawa, N. (2005). Mechanism of Nodal Flow: A Conserved Symmetry Breaking Event in Left-Right Axis Determination. Cell 121, 633–644.

Raya, A., Kawakami, Y., Rodríguez-Esteban, C., Ibañes, M., Rasskin-Gutman, D., Rodríguez-León, J., Büscher, D., Feijó, J. A. and Izpisúa Belmonte, J. C. (2004). Notch activity acts as a sensor for extracellular calcium during vertebrate left–right determination. Nature 427, 121–128.

Robinson, A. M., Takahashi, S., Brotslaw, E. J., Ahmad, A., Ferrer, E., Procissi, D., Richter, C.-P., Cheatham, M. A., Mitchell, B. J. and Zheng, J. (2020). CAMSAP3 facilitates basal body polarity and the formation of the central pair of microtubules in motile cilia. P. Natl. Acad. Sci. USA. 117, 13571–13579.

Schweickert, A., Weber, T., Beyer, T., Vick, P., Bogusch, S., Feistel, K. and Blum, M. (2007). Cilia-Driven Leftward Flow Determines Laterality in Xenopus. Curr. Biol. 17, 60–66.

Shinohara, K., Kawasumi, A., Takamatsu, A., Yoshiba, S., Botilde, Y., Motoyama, N., Reith, W., Durand, B., Shiratori, H. and Hamada, H. (2012). Two rotating cilia in the node cavity are sufficient to break left–right symmetry in the mouse embryo. Nat. Commun. 3:622.

Sorokin, S. (1962). CENTRIOLES AND THE FORMATION OF RUDIMENTARY CILIA BY FIBROBLASTS AND SMOOTH MUSCLE CELLS. J. Cell Biol. 15, 363–377.

Sulik, K., Dehart, D. B., Inagaki, T., Carson, J. L., Vrablic, T., Gesteland, K. and Schoenwolf, G. C. (1994). Morphogenesis of the Murine Node and Notochordal Plate. Dev. Dynam. 201, 260–278.

Supp, D. M., Witte, D. P., Potter, S. S. and Brueckner, M. (1997). Mutation of an axonemal dynein affects left–right asymmetry in *inversus viscerum* mice. Nature 389, 963–966.

Supp, D. M., Brueckner, M., Kuehn, M. R., Witte, D. P., Lowe, L. A., McGrath, J., Corrales, J. and Potter, S. S. (1999). Targeted deletion of the ATP binding domain of left-right dynein confirms its role in specifying development of left-right asymmetries. Development 126, 5495–5504.

Stephen, L. A., Johnson, E. J., Davis, G. M., McTeir, L., Pinkham, J., Jaberi, N. and Davey, M. G. (2014). The Chicken Left Right Organizer has Nonmotile Cilia which are Lost in a Stage-Dependent Manner in the talpid^3^ Ciliopathy. genesis 52, 600-613.

Takao, D., Nemoto, T., Abe, T., Kiyonari, H., Kajiura-Kobayashi, H., Shiratori, H. and Nonaka, S. (2013). Asymmetric distribution of dynamic calcium signals in the node of mouse embryo during left–right axis formation. Dev. Biol. 376, 23–30.

Takeda, S., Yonekawa, Y., Tanaka, Y., Okada, Y., Nonaka, S. and Hirokawa, N. (1999). Left-Right Asymmetry and Kinesin Superfamily Protein KIF3A: New Insights in Determination of Laterality and Mesoderm Induction by *kif3A*^-/-^ Mice Analysis. J. Cell Biol. 145, 825–836.

