## Supplementary material for "Left-right asymmetry is formed in the basal bodies of the mouse node cilia in a cilia motility-dependent manner": Fig. S1

### Supplementary information

Fig. S1

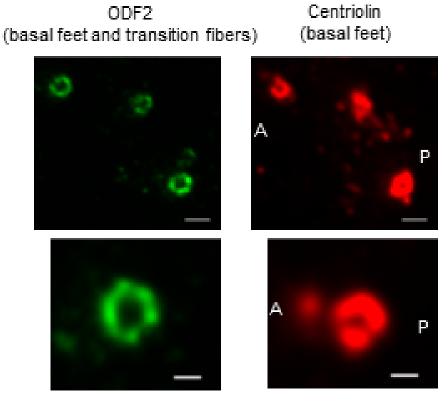

Fig. S1

**No L/R asymmetry was found in the arrangement of the basal feet around the basal bodies of the mouse node cilia.** XY-projected images (maximum intensity projection) of immunostaining for ODF2 and centriolin in the node of a somite-2 embryo. ODF2, which localizes to both basal feet and transition fibers, showed ring-like arrangements around the mother centriole as expected. Centriolin, which localizes to basal feet, also showed ring-like structure without apparent L/R asymmetry. Images were obtained using STED microscopy and deconvolution by Lightening (Leica). Bars, 500 nm for upper two panels, 200 nm for lower two panels.
